# Mathematical Model of Mechanosensing and Mechanically Induced Collective Motility of Cells on Planar Elastic Substrates

**DOI:** 10.1101/2022.12.19.520914

**Authors:** Riham K. Ahmed, Tamer Abdalrahman, Neil H. Davies, Fred Vermolen, Thomas Franz

## Abstract

Cells mechanically interact with their environment to sense, for example, topography, elasticity and mechanical cues from other cells. Mechano-sensing has profound effects on cellular behaviour, including motility. The current study aims to develop a mathematical model of cellular mechano-sensing on planar elastic substrates and demonstrate the model’s predictive capabilities for the motility of individual cells in a colony.

In the model, a cell is assumed to transmit an adhesion force, derived from a dynamic focal adhesion integrin density, that locally deforms a substrate, and to sense substrate deformation originating from neighbouring cells. The substrate deformation from multiple cells is expressed as total strain energy density with a spatially varying gradient. The magnitude and direction of the gradient at the cell location define the cell motion. Cell-substrate friction, partial motion randomness, and cell death and division are included.

The substrate deformation by a single cell and the motility of two cells are presented for several substrate elasticities and thicknesses. The collective motility of 25 cells on a uniform substrate mimicking the closure of a circular wound of 200 μm is predicted for deterministic and random motion. Cell motility on substrates with varying elasticity and thickness is explored for four cells and 15 cells, the latter again mimicking wound closure. Wound closure by 45 cells is used to demonstrate the simulation of cell death and division during migration.

The mathematical model can adequately simulate the mechanically-induced collective cell motility on planar elastic substrates. The model is suitable for extension to other cell and substrates shapes and the inclusion of chemotactic cues, offering the potential to complement *in vitro* and *in vivo* studies.

## 1. Introduction

Cell motility is a complex and periodic process underlying physiological development and diseases. Cell motility has been described for different cell types and environments (Shellard and Mayor 2020; Yamada and Sixt 2019; Wang et al. 2019; Matte et al. 2019; Tusan et al. 2018; Ladoux and Mège 2017). It is considered that the motility of all cell types occurs by similar molecular mechanisms. Cells interact mechanically with the substrate through focal adhesions and apply forces that deform the surrounding substrate area. Neighbouring cells sense these forces and deformations (Trepat et al. 2009; Throm Quinlan et al. 2011; Oakes 2018; Angelini et al. 2010; Gov 2009; Reinhart-King et al. 2008) and move in the direction of principal signals. Cells sense physical changes in the substrate, such as topographical traits and stiffness gradients (Hadjipanayi et al. 2009; Schwarz and Safran 2013). Cells can identify substrate rigidity based on the strength of a mechanical signal they sense (Lo et al. 2000; Yip et al. 2013). Changes in the cellular environment guide cell motility, e.g., cells move towards stiffer substrate regions (Lo et al. 2000; Gupton and Waterman-Storer 2006; Hadjipanayi et al. 2009).

Cell motility depends on various factors such as cell type, single or collective locomotion, substrate and extracellular matrix (ECM), adhesion strength, and organisation of the cellular cytoskeleton. Cells move individually and collectively, the latter particularly in wound healing (Liang et al. 2007; Arciero et al. 2011), tissue regeneration (Zorn et al. 2015; Angelini et al. 2011) and cancerous tumour invasion (Mendoz and Lim 2011; Rørth 2009). In a collective, cells interact with the substrate and neighbouring cells (Reinhart-King et al. 2008). The cell-substrate interaction can affect the rate and directionality of the collective cell migration (Liang et al. 2007; Borau et al. 2011; Bloom and Zaman 2014).

*In vitro* experiments have become more sophisticated and complex to generate accurate findings similar to *in vivo* studies. Mathematical models and computational simulations can generate reproducible and controlled experiments at significant lower costs. These models can be characterised according to different factors such as the numerical approach: continuous (Vermolen and Javierre 2012; Blanch-Mercader and Casademunt 2017), discrete (Szabó and Merks 2013; Marée et al. 2007; Charteris and Khain 2014), or hybrid (González-Valverde and García-Aznar 2018; Vittadello et al. 2018; Escribano et al. 2018). They can also be characterised according to the description of cell motility: individual (Dokukina and Gracheva 2010; Flaherty et al. 2007; Danuser et al. 2013; Gracheva and Othmer 2004; Allena et al. 2016; Dallon et al. 2013) or collective (Giniūnaitė et al. 2020; Satulovsky et al. 2008; Yogurtcu et al. 2012; Camley and Rappel 2017). Some of the models can be characterised as mechanical (Zaman et al. 2005; Borau et al. 2011; Mousavi et al. 2014a; van Oers et al. 2014), biochemical (Tan and Chiam 2018; Del Amo et al. 2017; Liebchen and Löwen 2018), or mechano-chemical (Fang et al. 2022; Löber et al. 2015; Marzban et al. 2019).

Recently, combinations of methodologies between computational modelling and *in vitro* experiments have opened new opportunities for researchers to directly obtain additional information of specific factors on individual and collective motility that can be indirectly evaluated from *in vitro* experiments to mention a few, (Kim et al. 2018; Staddon et al. 2022; Merino-Casallo et al. 2018; Mousavi et al. 2013).

Some computational models have incorporated stochastic processes such as cell proliferation (Mousavi and Doweidar 2016; Mousavi and Doweidar 2015) and cell death (Chen et al. 2020, 2018), and have used random walk to mimic cell migration (Safaeifard et al. 2018; Vermolen et al. 2015). However, most of these models did not consider the interactions between cells.

The mechanical communication of cells through substrate deformation due to traction forces has been considered previously in experimental and computational studies (Reinhart-King et al. 2008; Nerger et al. 2017; Franck et al. 2011; Maskarinec et al. 2009; Vermolen and Gefen 2012; Ben-David and Weihs 2021). These studies focused on capturing each cell’s motility on the substrate. Neighbouring cells on the substrate attract and move toward each other based on their traction forces. Cell locomotion, propagation, and apoptosis are considered with a semi-random probability distribution. Other models incorporate additionally cell-cell interactions in cell aggregates in the prediction of collective cell migration (Staddon et al. 2022; Marzban et al. 2019; Blanch-Mercader and Casademunt 2017; Löber et al. 2015; Peng and Vermolen 2020)

The current study aims to develop a mathematical cellular mechano-sensing model and demonstrate the predictive capabilities for the motility of individual cells in a colony on planar elastic substrates.

## 2. Methods

### 2.1 Mechanically induced cell motility

Two or more cells (of the same cell type) are considered attached to and exerting traction forces on a planar substrate. Considering traction forces perpendicular to the substrate surface is inspired by the experimental work of several groups (Nerger et al. 2017; Franck et al. 2011; Maskarinec et al. 2009; Ben-David and Weihs 2021; Reinhart-King et al. 2008). The traction forces deform the substrate underneath and in the vicinity of each cell. Each cell senses the deformation gradient induced in the substrate by neighbouring cells. For simplicity, the planar substrate is assumed as an elastic foundation attached to a rigid plate, and the cells are assumed to be rigid disks (Figure 1).

**Figure 1.**
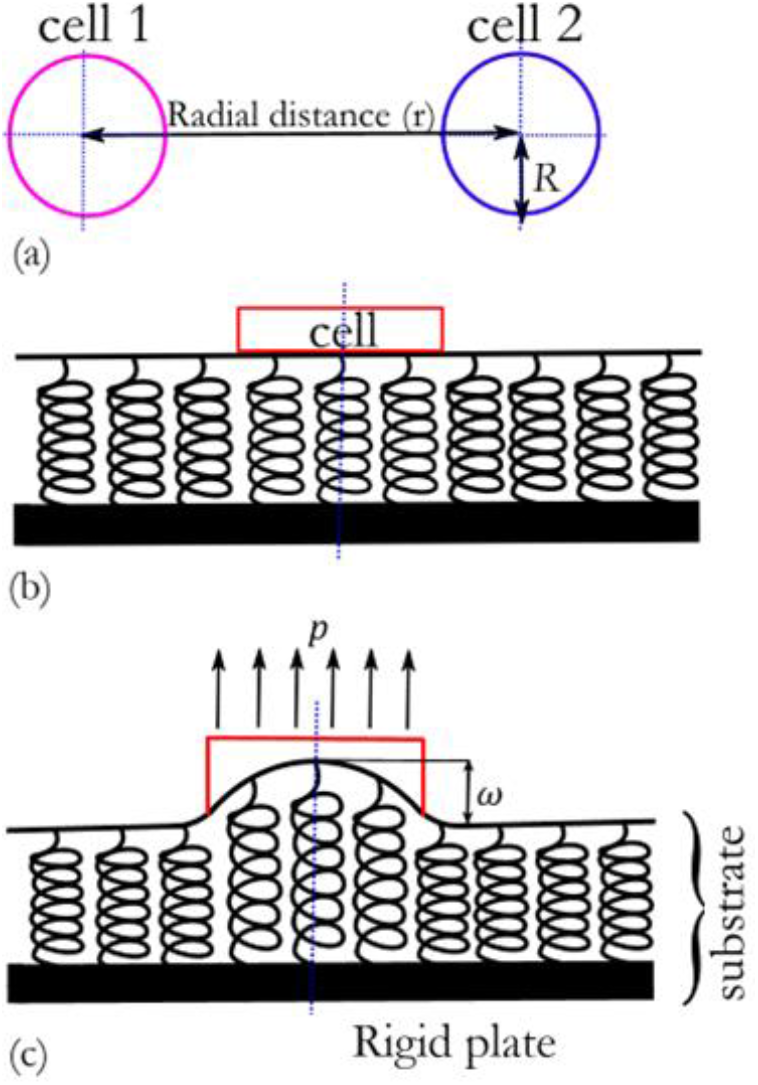
Schematic illustration of cells, planar substrate, and cell-substrate interaction. a) Top view of two cells with radius *R* at a distance *r* between their centres. b) Side view of a single cell (red) on an elastic substrate (black), represented by spring elements on a rigid plate. c) Deformation of the elastic substrate underneath and around a single cell due to cellular traction force *p* exerted on the substrate.

With ***c**_i_*(*t*) = (*x_i_*(*t*), *y_i_*(*t*)) as the centre position of the cell *i* at a time *t*, the substrate deformation, *ω*, underneath and around each cell *i* (Loof 1965) is:

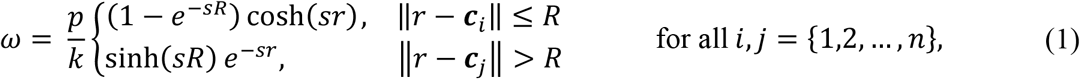

where *R* is the cell’s radius, *p* is the traction force distributed on the substrate over a cell area, *r* is the distance between the centres of two cells, ***c**_j_*(*t*) denotes the centre positions of cells *j* in the neighbourhood of cell *i*, and

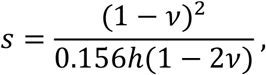

with *v* as the Poisson’s ratio and *h* as the thickness of the substrate. For a uniformly distributed force, the substrate stiffness, *k*, (Loof 1965) is:

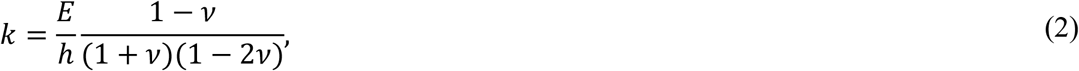

where *E* denotes the elastic modulus of the substrate.

With the strain energy obtained from the extension of the Pasternak foundation model (Selvadurai 1979), the strain energy density (i.e., strain energy per unit volume) can be expressed as a function of the substrate deformation:

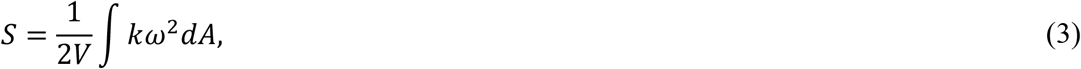

where *V* is the deformed volume of the substrate.

It is assumed that each cell is a source and a receiver of strain energy density signals due to cellular traction forces on the substrate. The total strain energy density, *ψ*, of each cell *i* is the strain energy density due to the substrate deformation induced by cell *i*, and the strain energy densities from other cells *j* that cell *i* senses:

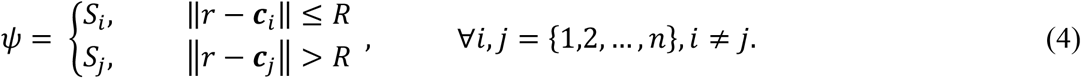

Since the strain energy density is additive, Eqn. (4) can be rewritten as:

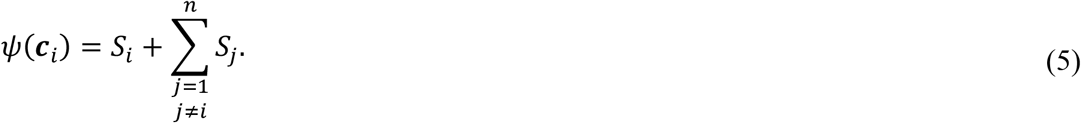

Equation (5) represents the sum of all strain energies. The first term represents the strain energy density under cell *i* based on the substrate deformation due to its traction force. The second term represents the sum of all strain energy densities under the same cell *i* resulting from substrate deformation caused by traction forces of other cells in the domain.

The migration direction of each cell *i* is the direction of the strain energy density resultant:

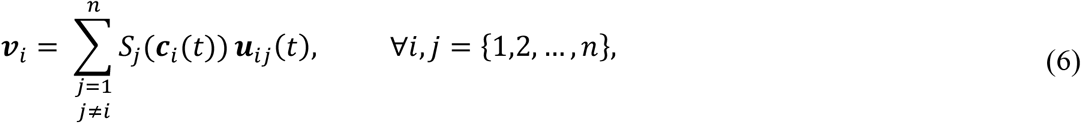

where ***u**_ij_*(*t*) denotes the unit vector connecting cell *i* to other cells *j*, i.e.,

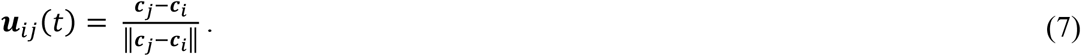

The normalised form of the motility unit vector is:

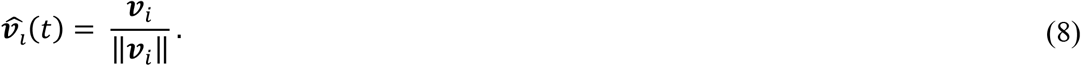

The magnitude of the displacement is assumed to be proportional to the strength of the mechanical signal, i.e., the substrate deformation. Thus, the movement of cells towards each other is:

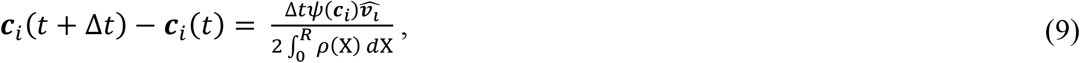

Where *ρ*(X) denotes the friction coefficient of the cell-substrate adhesion, and X denotes the position of a point in the cell that varies in the interval occupied by the cell, i.e., X = [−*R*, *R*],

### 2.2 Cell-substrate interaction

Individual cells are considered homogenous rigid components. Suppose that the effective friction coefficient of the cell-substrate adhesion, *ρ*(X), is linearly dependent on the concentration of integrins attached to substrate ligands. Therefore, the cell adhesion strength is:

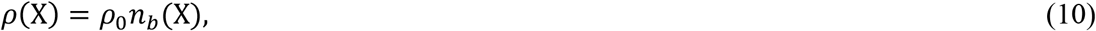

where *ρ*_0_ is the friction coefficient for a cell receptor binding to the substrate, and *n_b_*(X) is the density of bound integrins.

Suppose *n_f_* is the concentration of free integrins and *n_s_* is the concentration of the substrate ligands. The propagation of free integrins *n_f_*, the attachment of integrin receptors to substrate ligands and the detachment of integrin receptors from substrate ligands are assumed as instantaneous mechanisms compared to the time scale of cell motility. It is also assumed that the bond formation rate *k_f_* is constant, the dissociation rate 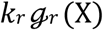 varies with the position in a cell, and the density of substrate ligands *n_s_* is uniform and constant. Ligand depletion can be neglected when the ligand number is saturated. Then, the concentration of free integrins *n_f_* can be considered constant, and the density of the integrins is:

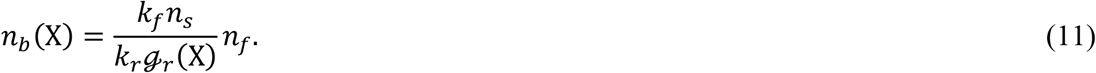

To represent the cell adhesion dynamics, suppose that:

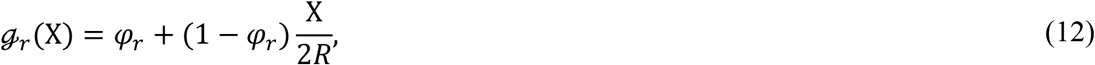

where *φ_r_* ≥ 1 denotes a decrement of the dissociation rate from the rear to the front of the cell.

The total number of integrin receptors is assumed to be constant over time and obtained from Gracheva and Othmer (2004) and Zaman et al. (2005). The concentration of integrin receptors and free integrins *n_f_* are estimated from Dai et al. (2015). With parameter *k_s_* = *k_f_ n_s_/k_r_* representing the cell-substrate interaction, the friction coefficient of cell-substrate adhesion is:

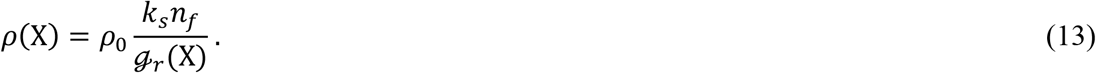

### 2.3 Stochastic processes in cell motility

#### 2.3.1 Cell motility with randomness

Cell motility is a biological and partial random process. Thus, a certain degree of motion randomness is incorporated into the present model using the Brownian motion approach. The standard Brownian vector, Δ***B***(*t*), has two independent entries that are normally distributed with zero mean and variance Δ*t*, that is, the entries of Δ***B***(*t*)~*N*(0, Δ*t*). Hence, for cells with partial random motion:

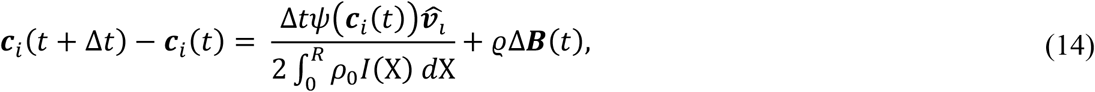

where *ϱ* considers probabilistic variations due to uncertainties, such as tissue composition, cell composition, and access to necessary chemicals. *ϱ*Δ***B***(*t*) is commonly referred to as random walk. The definition of Brownian motion can be found, for instance, in Karatzas and Shreve (2012) and (Bressloff 2014).

#### 2.3.2 Cell death and division

Cell death and division can be represented as random processes since cell cycles differ from cell to cell (Kar et al. 2009). Here, cell death and division are modelled using stochastic principles described in Chen et al. (2018) and Chen et al. (2020). For simplicity, each cell in the population is assumed to be either viable or dead, purposely disregarding that a cell may mutate or be injured and then recover or die. The probability *P_i_* of a cellular event of cell *i*, i.e., death or division, within a time interval Δ*t* is considered to follow an exponential distribution and calculated by:

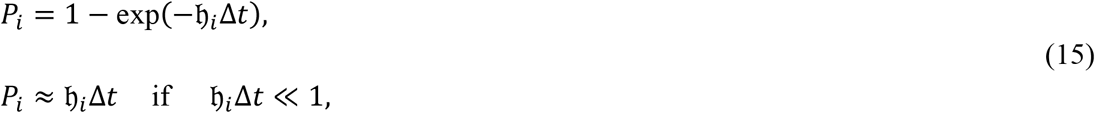

where 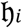 is the probability rate of cellular event per second. At each time interval Δ*t*, a cellular event 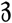 is generated from the standard uniform distribution U(0, 1), i.e., 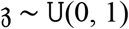, such that:

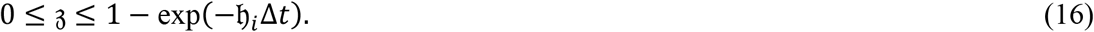

In this model, cell death and division are simulated using the probability rates 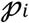 and 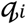, respectively. The probability rate for the death of cell *i* is assumed to be 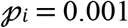 (Vermolen et al. 2012). The probability rate of cell death is generally related to the cell’s condition or environment, such as temperature and pH level. The probability rate that cell *i* dies within a time interval Δ*t* does not depend on the size of time step of the numerical solution.

In this study, the degradation of dead cells is ignored, and since only mechanical effects are considered, the chemical effect of dead cells is neglected. Moreover, any influence of dead cells on the motility of viable cells is ignored. It is further assumed that a cell’s death is not affected by its environment and is sudden and discrete. Dead cells are removed from the simulation immediately at each time frame Δ*t*, and their vacated positions can be occupied by any viable cell.

The same probabilistic principles are applied to cell division. Let 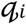 be the division probability rate of each cell *i* during a time interval Δ*t*. The probability rate for cell *i* to divide is assumed to be 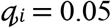 (Vermolen et al. 2012). For the division of a cell into two cells, the original cell is shifted by a distance *R* in a random direction and a new daughter cell is added to the model such that their edges resemble the centre position of the original cell. It is assumed that once the original cell and the daughter cell are in contact, they stay together until the motility force changes their polarisation orientation, after which they may separate. This is a simplifying essential assumption; hence, the authentic tendency is also a stochastic parameter.

### 2.4 Cell motility simulations

The computational domain of the planar substrate is considered as 1,000 μm × 1,000 μm for all cell motility simulations except those involving cell death and division, for which 1,500 μm × 1,500 μm is used. The cells with a radius *R* = 50 *μ*m are considered. The simulations mimic hypothetic cases; hence, hypothetic values have been used for many input parameters. The values of the model parameters introduced in the above equations are listed in Table 1 unless stated otherwise.

**Table 1.**
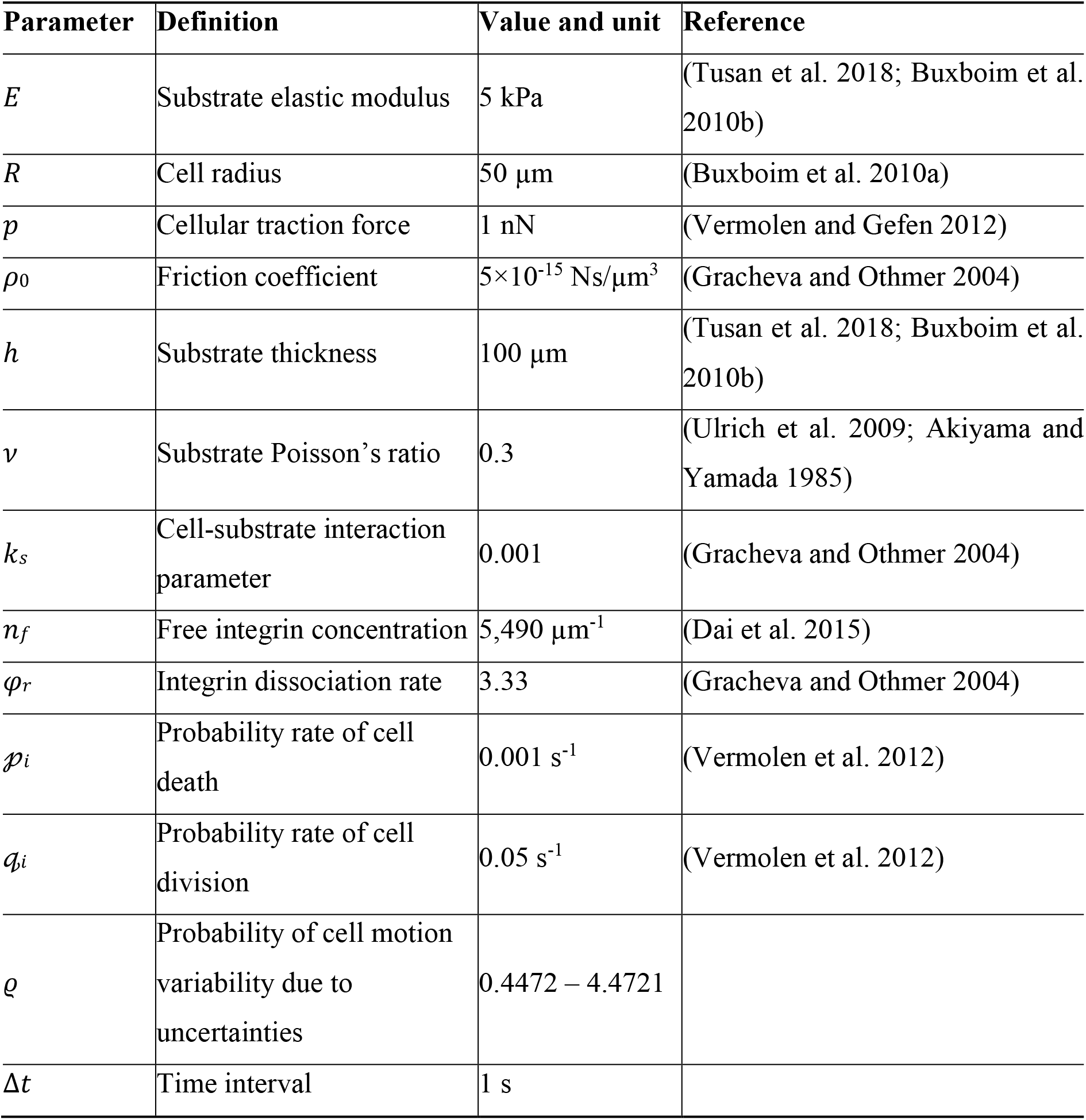
Parameters and values used in the model.

Each simulation experiment runs with a time step of 10 s. It is worth noting that the time scale varied and depended on the migration velocity. Of course, there is a likelihood of a small displacement vector, i.e., close to zero, during one or more time steps; hence a cell may not have migrated to a new location, and the cell velocity may be close to zero during these time steps.

First, the evolution of the 2D substrate deformation underneath a single cell is investigated for different values of the elastic modulus (*E*) and thickness (*h*) of the planar substrate. After that, the motility due to cellular mechanical interactions and the effect of substrate elasticity and thickness are studied. Finally, the model’s capability of incorporating cell death and division in collective cell motility is demonstrated.

Cell deformation due to cell-cell contact (Peng and Vermolen 2020) is considered in the model by permitting cells to overlap up to 30%, after which repulsing of the cells is assumed. This condition is implemented by a negative attraction vector in Eq. (7) that corresponds to the strain energy density for 30% cell deformation This penalty condition prevents 100% overlap of the cells.

The mathematical model has been implemented in MATLAB (The MathWorks, Inc, Natick, MA, USA). The custom code files are available for download as indicated in the Data availability section.

## 3. Results

### 3.1 Substrate deformation by a single cell

The deformation of the substrate normal to its surface due to a uniformly distributed traction force *p* of a cell with radius *R* = 50 *μ*m is determined for different values of the substrate elastic modulus and thickness (Figure 2). The substrate deformation decays exponentially with distance from the cell centre, and it increases with decreasing elastic modulus and increasing thickness of the substrate.

**Figure 2.**
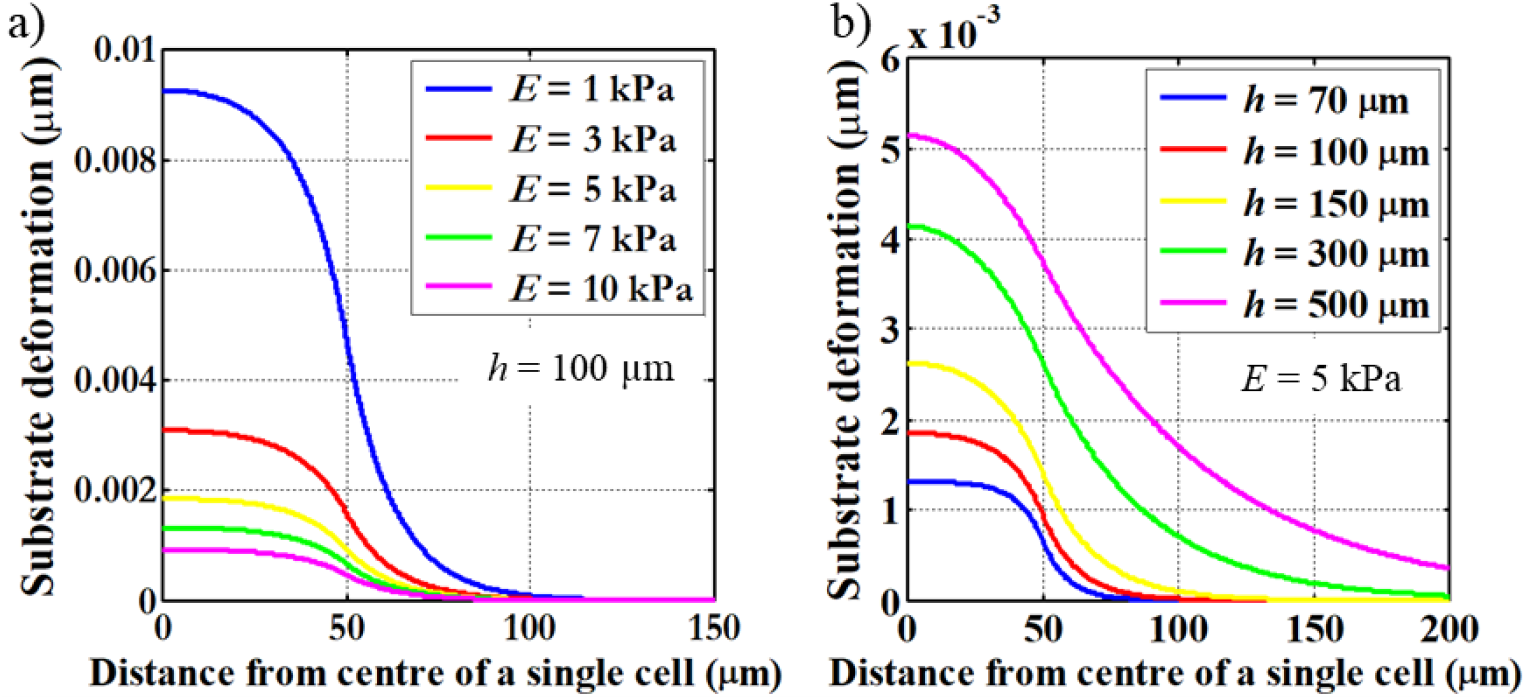
Substrate deformation by the distributed traction force *p* of a single cell with radius R = 50 *μ*m versus distance from cell centre for different values of the substrate elastic modulus *E* with constant substrate thickness *h* = 100 μm (a) and thickness *h* with constant *E* = 5 kPa (b). The substrate deformation normal to the substrate plane decays exponentially from the cell centre, and the magnitude of the substrate deformation increases with decreasing substrate elasticity *E* (a) and increasing substrate thickness *h* (b).

### 3.2 Effect of substrate elasticity and thickness on the migration velocity of two approaching cells

The movement of two cells with fully deterministic motion towards each other on a substrate is considered for different substrate elastic modulus and thickness values. The migration velocity of the cells increases with decreasing substrate elasticity (Figure 3a) and thickness (Figure 3b). The initial distance between the two cells is random, i.e., different for each case (variation of elastic modulus and thickness, respectively) but the same for the parametric simulations of each case. Cell viability is defined as a discrete parameter, and continued cell viability is ignored, implying that a cell is either fully viable or dead.

**Figure 3.**
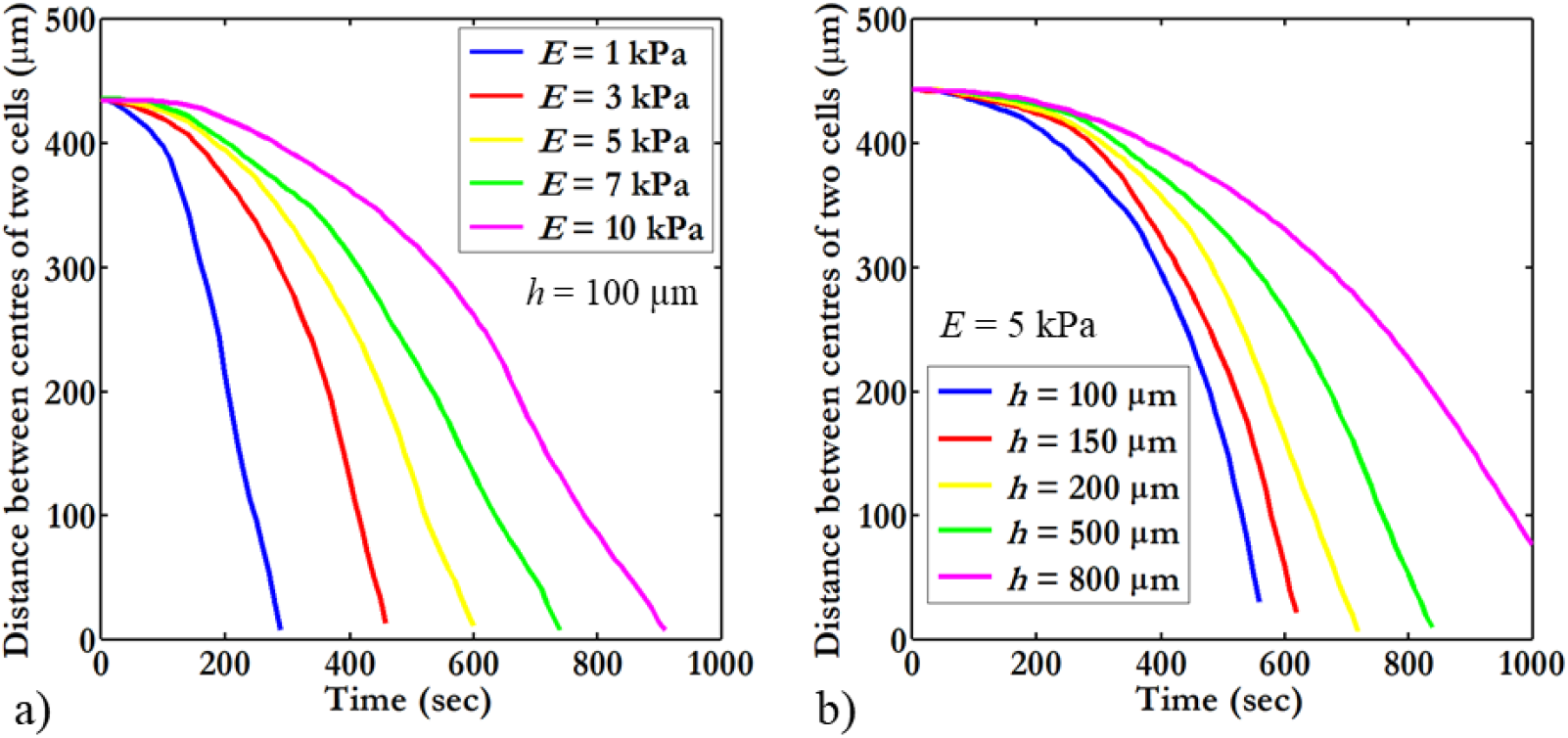
Distance versus time of two cells approaching each other in fully deterministic motion (i.e., absence of stochastic motion) on a planar substrate for different substrate elasticity and thickness. The migration velocity of the cells increases with decreasing substrate elasticity *E* (a, for *h* = 100 *μ*m) and substrate thickness (b, for *E* = 5 kPa).

The final distance between the two cells is smaller than the cell diameter (2*R* = 100 μm), i.e., the cells impinged on each other since repulsive forces are not considered.

### 3.3 Collective cell motility mimicking wound closure

Collective cell motility due to mechanical cues on a planar substrate with uniform elastic modulus and thickness with and without standard Brownian motion (random movement) is investigated using 25 cells arranged randomly in two concentric rows of cells (with an outer radius of 400 *μ*m). The arrangement provides a central cell-free space mimicking a wound with a radius of approximately 200 μm. The probabilities of cell death 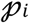 and cell division 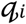 are assigned small values to emphasise the mechanically-induced cellular motion, i.e., 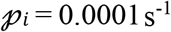 and 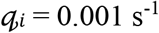.

With fully deterministic motion (Figure 4a), cells initially move towards neighbouring cells in the dominant direction of the resultant strain energy density gradient in the substrate (*t* = 22 s). In addition, the cells in contact deform (shown as overlap) as inter-cellular forces are not considered in the model. The interaction between cells mainly depends on the initial cell positions. Once cells are in contact, they start migrating together into the central cell-free area (*t* = 44 s). The cells fill the cell-free central area following further migration towards the centre regardless of the initial cell locations (*t* = 66 s). Including stochastic perturbations in the model (Figure 4b) adds variability in the direction and magnitude of the cellular motion during each solution time step (*t* = 0, 20, and 40 s). However, the stochastic perturbation does not change the final cluster arrangement (*t* = 61 s).

**Figure 4.**
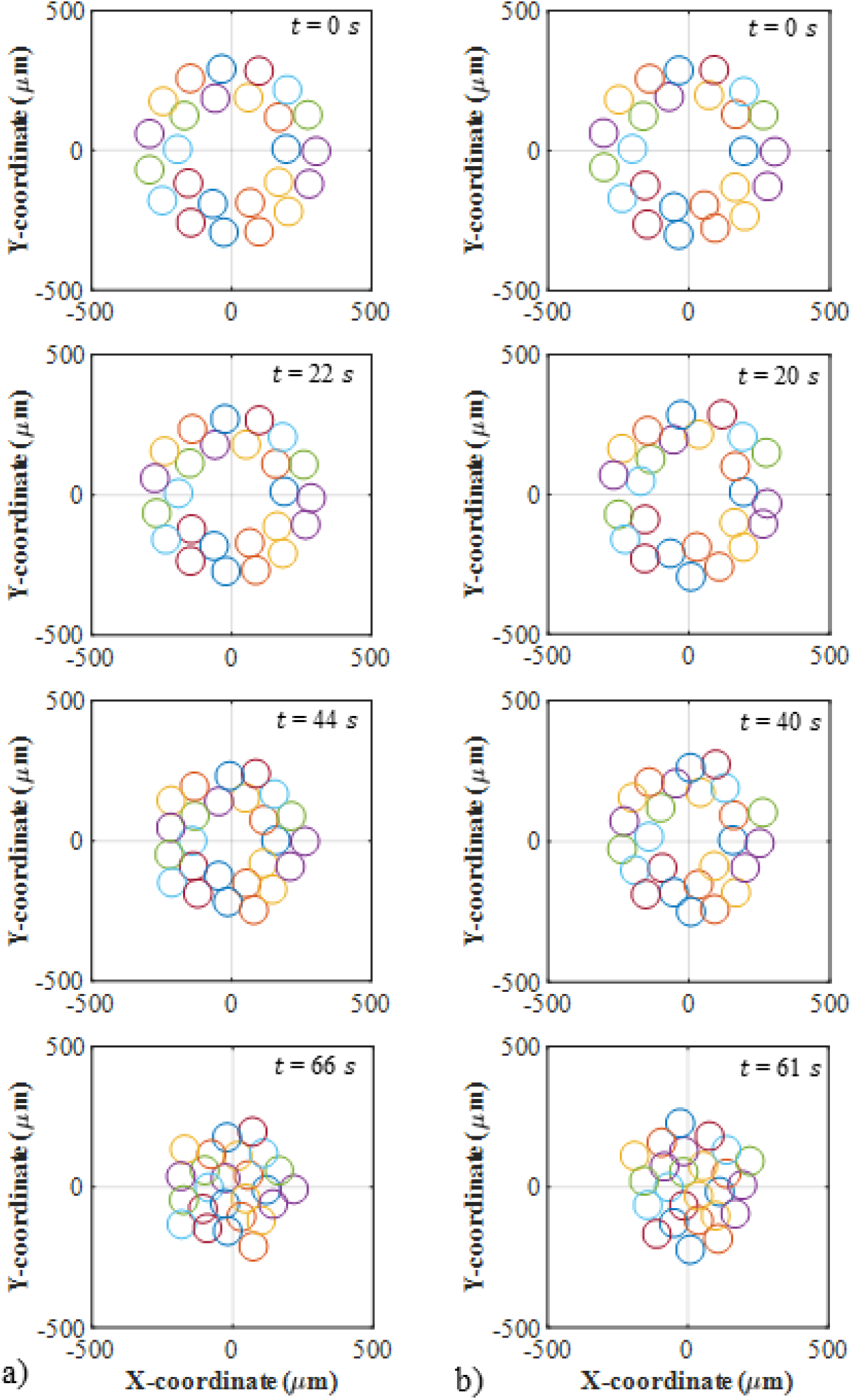
Positions of 25 cells on a planar substrate during collective migration with (a) fully deterministic and (b) randomly perturbed motion as a function of time. a) Initial positions of the cells at *t* = 0 s in two concentric rows (outer radius: 400 μm, inner radius: 200 μm) with a cell-free centre area. At *t* = 22 s, neighbouring cells move towards each other, directed by the resultant strain energy density gradient (see also supplemental video V4A). The cells contact each other and start deforming (represented by overlap) in the absence of repulsive intercellular forces. At *t* = 44 s, cells start forming clusters and moving into the cell-free central area (wound). At *t* = 66 s, cells collectively move into the centre and fill the entire cell-free central area. b) Initial positions of the cells at *t* = 0 s. At *t* = 20 s, neighbouring cells move towards each other following the resultant strain energy density gradient. At *t* = 40 s, cells start to form clusters and move toward the cell-free centre of the substrate. At *t* = 61 s, cells migrate into and close the cell-free central substrate area (see also supplemental video V4B). (Model parameter values are listed in Table 1 except 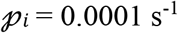 and 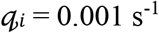.)

The evolution of locomotion paths of 10 cells arranged in a circle with fully deterministic and randomly perturbed motion illustrates the effect of Brownian motion, i.e., the deflection of the mechanically-induced motion (see Figure 5).

**Figure 5.**
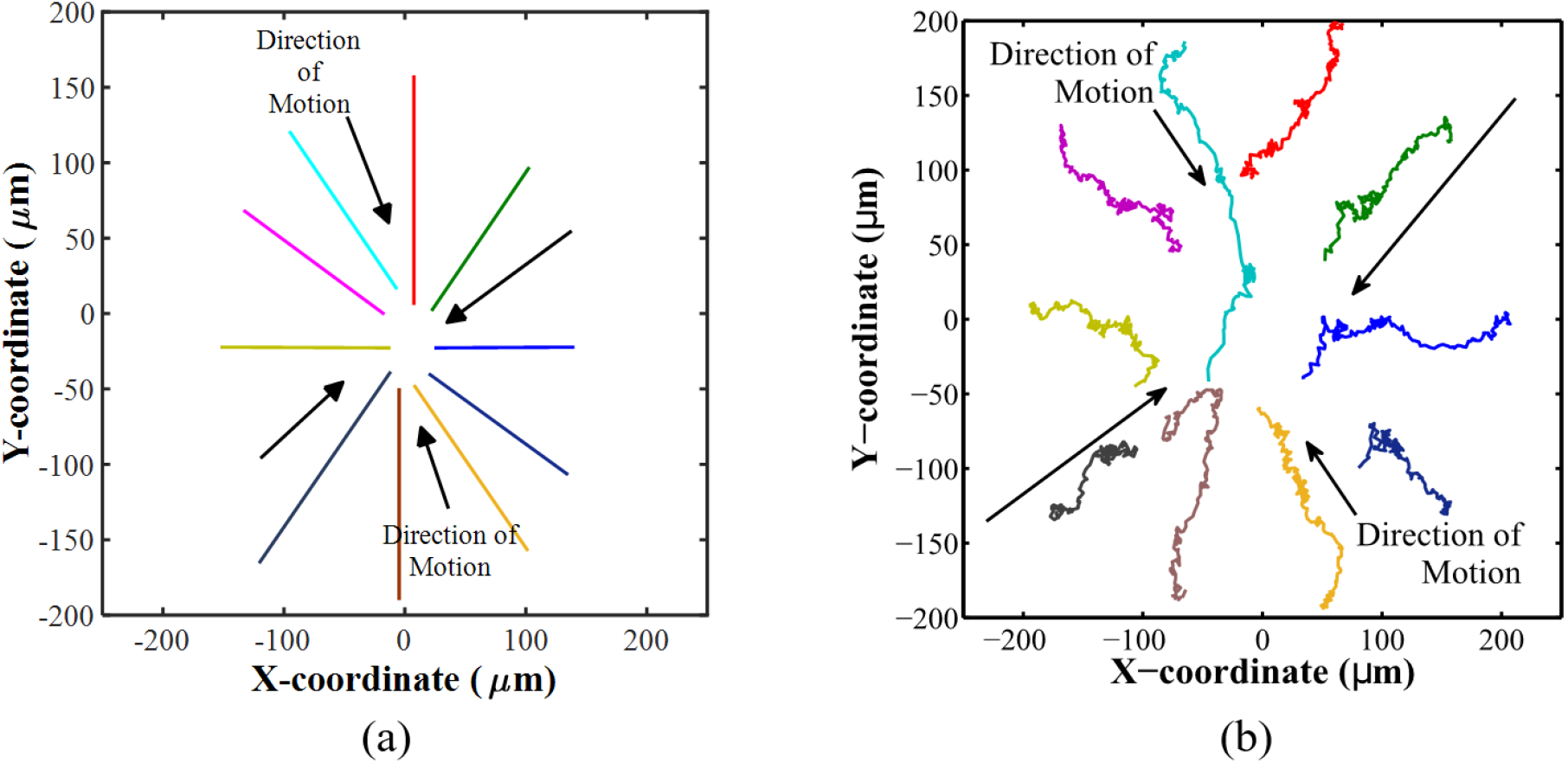
Trajectories of ten cells during collective motility for (a) fully deterministic and (b) randomly perturbed motion starting from an initial circular arrangement and moving for the same period. Coloured lines illustrate the movement paths of cell centre points. Black arrows indicate the overall direction of the cellular motion. For fully deterministic motion, cells move along straight paths towards the centre of the substrate (computational domain) and form a cluster. For perturbed motion, the trajectories show the randomness overlaying the mechanically induced motion of the cells. (Model parameter values are listed in Table 1 except 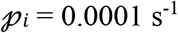 and 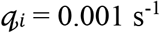.)

### 3.4 Cell motility for spatially varying substrate elasticity

To study the effect of substrate regions with different stiffness on cell motility, a planar substrate of 1,000 × 1,000 *μ*m with a constant thickness of *h* = 100 *μ*m is divided into a stiff (*E* = 15 kPa) and soft region (*E* = 5 kPa). The elastic modulus is changed as a step at X = 0 *μ*m, providing two regions with the dimensions of 500 × 1,000 *μ*m (Figure 6). Cell death and division are not considered; hence 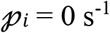 and 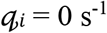 are used.

**Figure 6.**
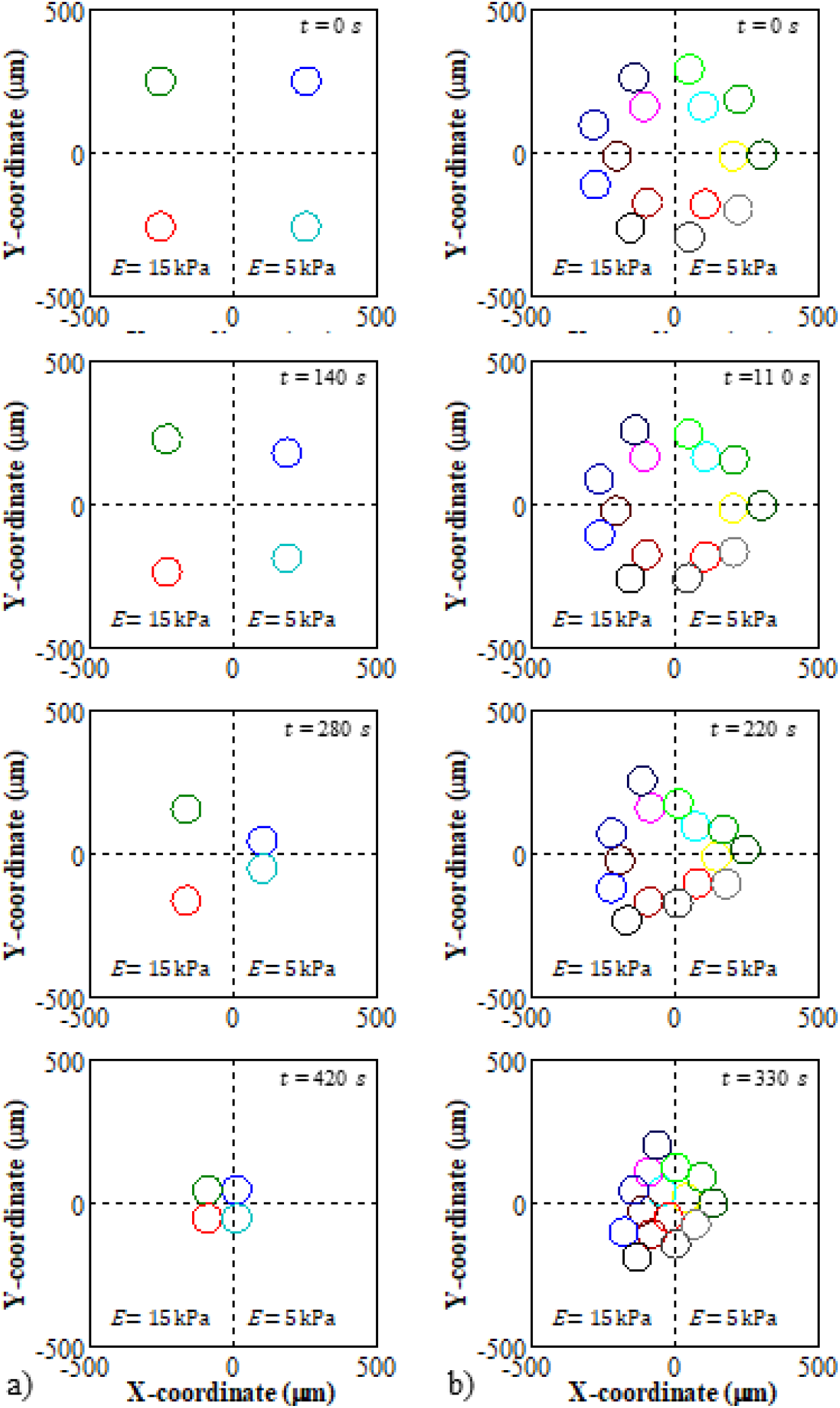
Positions of 4 (a) and 15 (b) cells migrating with fully deterministic motion on a planar substrate with varying elastic modulus *E* and constant thickness *h* = 100 *μ*m at as a function of time. *E* changes at X = 0 *μ*m from *E* = 15 kPa in the substrate region with X < 0 (left, stiff region) to *E* = 5 kPa in the region with X ≥ 0 (right, soft region). a) Initial positions of four cells at *t* = 0 s, with two cells each on the stiff and soft substrate region. After *t* = 150 s, the two cells in the left and right region, respectively, move towards each other (i.e., towards Y = 0) due to the mechanical cues. The cells migrate faster in the soft than the stiff region. At *t* = 300 s, the cells on the soft region reach each other (at Y ≈ 0) and continue moving collectively in the X-direction towards the cells in the stiff substrate region (see also supplemental Video V6A). At *t* = 515 s, the four cells cluster in the substrate centre, slightly towards the stiff substrate region. b) Initial ring-shaped arrangement of 15 cells at *t* = 0 s. At *t* = 150 s, the cells migrate towards each other on both stiff and soft substrate regions. At *t* = 235 s, the cells migrate farther and have larger velocities on the soft than the stiff region. At *t* = 330 s, the cells from both regions form a single cluster. Cells in the soft substrate region migrate more towards and into the stiff region, whereas cells in the stiff region migrate less towards the soft region (see also supplemental Video V6B). (Model parameter values are listed in Table 1 except for values provided above in this caption.)

The first case considers four cells, i.e., one cell in each substrate quadrant, with an equal distance between the two cells in the stiff and soft substrate region (Figure 6a). The two cells in each stiffness region of the substrate primarily move towards each other (i.e., towards Y = 0) with a minor movement component towards X = 0. The cells in the soft region move faster than in the stiff region. Once the two cells in the soft region reach each other (at Y ≈ 0), they move collectively in the X-direction towards the cells in the stiff region. Similarly, once the cells in the stiff region reach each other, they move collectively towards the cells in the soft region. The four cells finally form a cluster in the centre of the substrate, although slightly in the stiff region (see also supplemental Video V7).

For 15 cells in the ring-shaped arrangement (Figure 6b) on the same substrate as in the previous case, the cells initially move towards and connect with neighbouring cells. The cells in the stiff substrate region remain distributed in two- and three-cell clusters compared to the more connected cells in the soft substrate region (*t* = 150 s). The cell cluster in the soft substrate region joins two clusters in the stiff substrate region (*t* = 235 s). In the final configuration, all cells cluster slightly towards the stiff region in the substrate centre and fill the cell-free space (*t* = 330 s) (see also supplemental Video V8). The 15 cells fill the cell-free central substrate area after *t* ≈ 330 s, whereas this takes *t* ≈ 515 s for the four cells, indicating an increase in migration velocity with an increase in cell number.

### 3.5 Cell motility for spatially varying substrate thickness

Four cells are considered on a substrate with a step-wise change of thickness *h* along X = 0, from *h* = 300 *μ*m for X < 0 to *h* = 100 *μ*m for X ≥ 0, and a constant elastic modulus of E = 5 kPa. Initially, two cells are located in the thick and thin regions, and the cells in each region have the same distance from each other (Figure 7a, *t* = 0 s). Cell death and division are not considered here, and 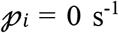 and 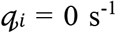. Initially, the single cells preferentially move towards each other, faster in the thin than the thick region (*t* = 106 s), until they are in contact at Y ≈ 0 in each region (*t* = 213 s). After that, the two two-cell clusters move towards each other and aggregate on the boundary between thin and thick substrate regions (*t* = 320 s). The cells keep moving around each other randomly and do not cross from the thick region to the thin region of the substrate (see also supplemental Video V7A).

**Figure 7.**
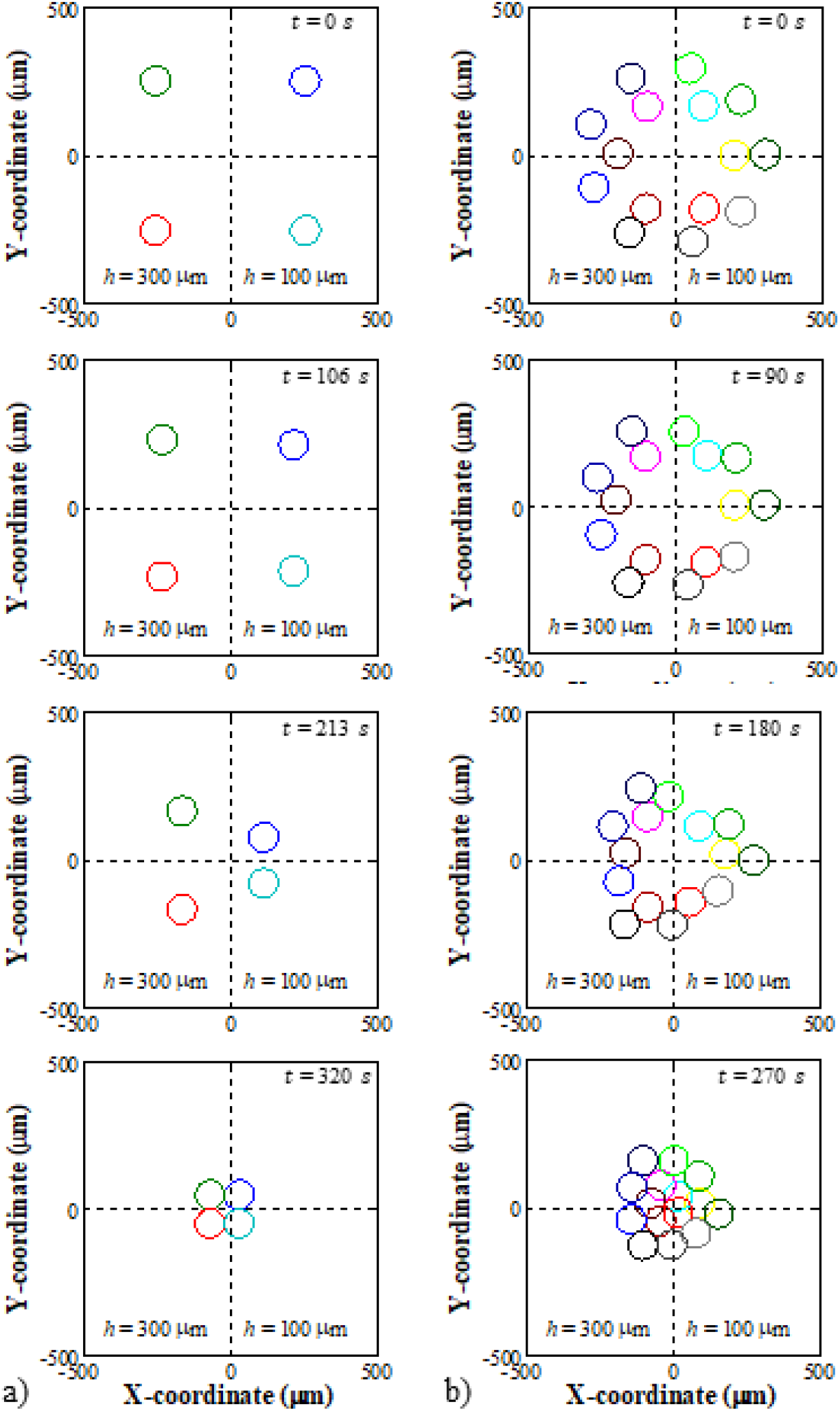
Positions of cells migrating with fully deterministic motion on a substrate with a step-wise change of thickness *h* along X = 0, from *h* = 300 *μ*m for X < 0 to *h* = 100 *μ*m for X ≥ 0, and a constant elastic modulus of *E* = 5 kPa. a) Initial configuration of four cells with the same distance between the two cells in the thin and thick substrate region at *t* = 0 s. At *t* = 106 s, the two cells in each substrate region preferentially move towards each other. Cells in the thin region migrate at a higher rate. At *t* = 213 s, the two cells in each region join at Y ≈ 0. At *t* = 320 s, the two two-cell clusters move towards each other and aggregate at X ≈ 0 slightly towards the thick substrate region (see also supplemental Video V7A). b) The initial circular arrangement of 15 cells with a cell-free central substrate area at *t* = 0 s. At *t* = 90 s, cells move to neighbouring cells. Cells in the thin substrate region but close to the thickness interface at X = 0 *μ*m move towards and into the thick substrate region. At *t* =180 s, the cells move collectively towards the substrate centre into the cell-free region. At *t* = 270 s, the cells fill the cell-free central substrate area (see also supplemental Video V7B). (Model parameter values are listed in Table 1 except for values provided above in the caption.)

The collective motility of 15 cells (Figure 7b) from an initial circular arrangement on a substrate with two thickness regions (as in the previous case) starts with the movement of individual cells towards neighbouring cells. However, cells on the thin substrate region near the thickness change moved towards and into the thick substrate region, but not vice versa. The overall migration tendency of cells is towards the thick substrate region (*t* = 90 s). The cells move collectively towards and into the cell-free central substrate region, with similar migration speeds in the thick and thin substrate regions (*t* = 180 s). Finally, the cells fill the central cell-free substrate area and cluster slightly more in the thick substrate region (*t* = 270 s) (see also supplemental Video V7B).

### 3.6 Cell motility with cell death and division

The prediction of collective cell motility with cell death and division is demonstrated with 45 cells in two concentric rows with an inner and outer radius of 400 *μ*m and 550 *μ*m (Hettler et al. 2013; Lan et al. 2010), respectively, on a uniform substrate with an elastic modulus of *E* = 5 kPa and a thickness of *h* = 100 *μ*m. The probabilities of cell death and division are set to 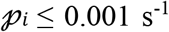 and 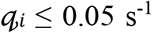 (Vermolen et al. 2012) as required for the different cases, i.e., collective motility with (i) cell death only, (ii) cell division only, and (iii) cell death and division. The larger number of 45 cells for these simulations compared to 15 and 25 cells in the previous cases are chosen based on the larger initial circular cell configuration.

For the case of motility with cell death only, the number of cells decreases during the collective migration into the central, cell-free substrate area from initially 45 cells to 40 cells after *t* = 115 s when the central wound area is closed (Figure 8a). For motility with cell division only, the cell number increases from 45 to 72 while the cells migrate into and close the central cell-free substrate area in 50 s (Figure 8b). For collective cell motility with cell death and division, the central cell-free substrate area is closed after *t* = 60 s while the cell number increases from 45 to 66 (Figure 8c).

**Figure 8.**
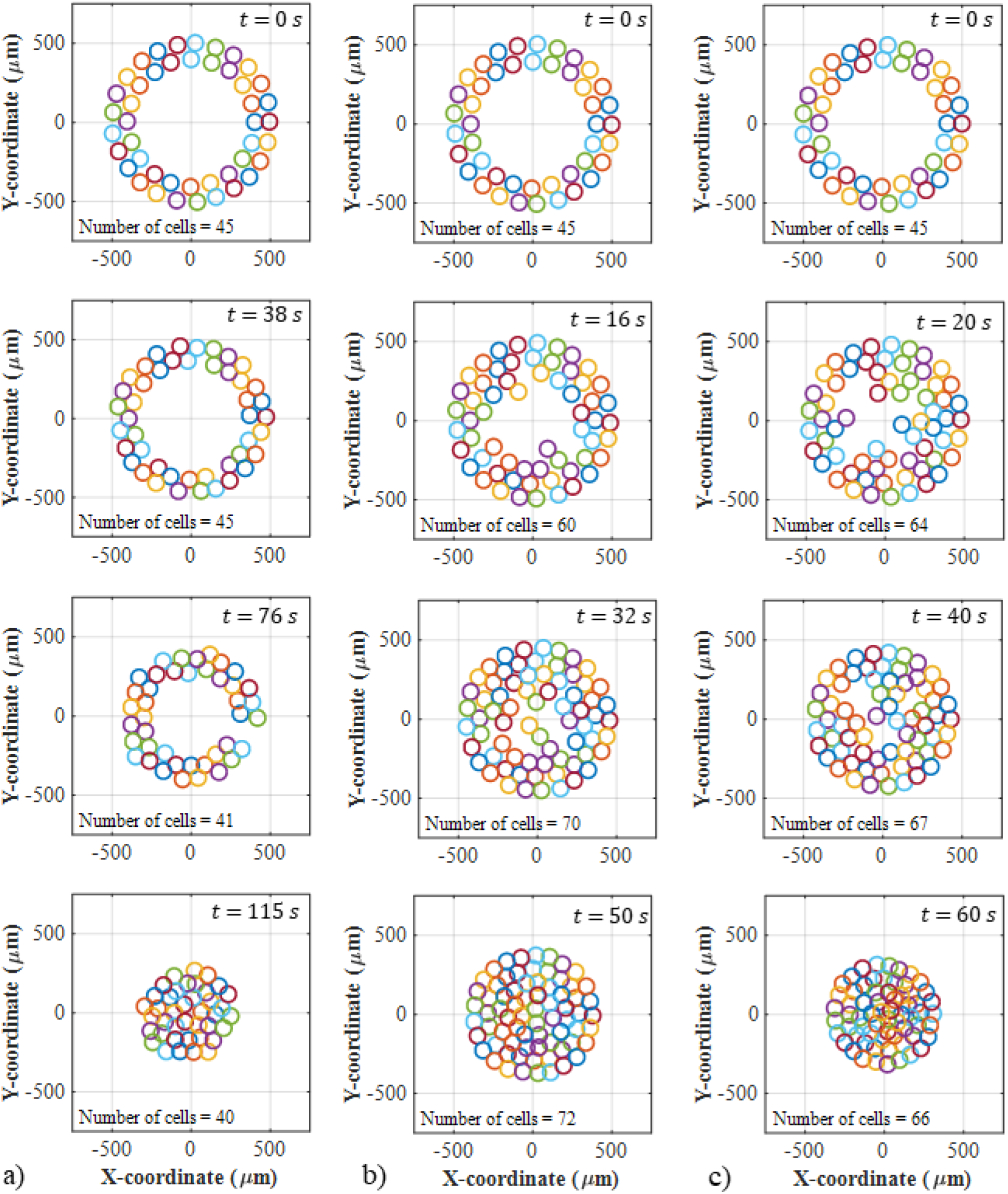
Positions of initially 45 cells on a planar substrate and changes in number and position of cells over time during collective migration with fully deterministic motion with (a) cell death only, (b) cell division only, and (c) cell death and division The initial arrangement of 45 cells in two concentric rows (inner radius: 400 μm, outer radius: 550 μm) is the same for the three cases (at *t* = 0 s). a) For cell death only, the cells close the cell-free central substrate area in *t* = 115 s and the number of cells decreases from 45 to 40. b) For cell division only, the central cell-free substrate area is closed in *t* = 50 s, and the cell number increases from 45 to 72. c) For cell death and division, the central cell-free substrate area is closed after *t* = 60 s, and the cell number increases from 45 to 66. (Please also see supplemental videos V8A to V8C for cases a to c. The supplemental video V8D shows cell motility with randomly perturbed motion with cell death and division not included above.) (Model parameter values are listed in Table 1 except for cell death and division probabilities as follows: a) 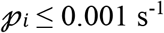, 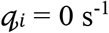, b) 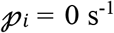, 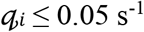, c) 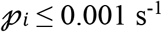, 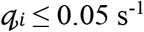.)

## 4. Discussion

A two-dimensional (2D) model for the simulation of mechanically induced motility of cells on a planar substrate is developed. The chosen approach is different from those proposed in previous studies (Zorn et al. 2015; González-Valverde and García-Aznar 2018; George et al. 2017; Camley and Rappel 2017; Kim et al. 2016) in considering the mechanical properties of the cellular microenvironment and cell-substrate adhesion with the initial goal of analysing how mechanical changes of the substrate guide cell migration.

### 4.1 Single-cell substrate deformation and migration velocity for varying substrate elasticity and thickness

The developed model indicates that substrate elasticity and thickness are essential in cell migration. An increase in substrate deformation due to traction forces of a single cell is observed for decreasing stiffness and increasing thickness of the substrate (Figure 2). The predicted effect of substrate stiffness is consistent with reports (Yip et al. 2013; Voloshin 2016; Gallinaro et al. 2013). The increasing deformation restriction may explain the increase in substrate deformation with the increase in substrate thickness in an elastic layer with decreasing thickness connected to a rigid foundation. Similar changes in substrate mechanics with substrate thickness are observed experimentally for several cell types (Buxboim et al. 2010a; Buxboim et al. 2010b; Merkel et al. 2007).

The increasing migration velocity of two interacting cells predicted for a decreasing elastic modulus, i.e., softening, of the substrate (Figure 3a) correlates with the increasing substrate deformation (i.e., mechanotactic cue) by a single cell for decreasing substrate elastic modulus (Figure 2a) and agrees with experimental reports (Lo et al. 2000; Yip et al. 2013; Adlerz et al. 2016). However, the increase in migration velocity of two cells with a decrease in substrate thickness (Figure 3b) contrasts the decrease in substrate deformation with a decrease in substrate thickness for a single cell (Figure 2b). Increased mechanosensing of cell colonies compared to single cells (Tusan et al. 2018) and the slowing down of cells with increasing substrate strain (van Oers et al. 2014) may explain the model prediction, at least in part.

### 4.2 Collective cell motility on uniform substrates

The model results reveal several differences between collective and single-cell migration on a planar substrate. Compared to the migration of two cells, the mechanical interactions between cells in colonies (i) delay their arrival at the final destination and (ii) increase the magnitude of the strain energy density and local velocity, which is also reported by Mousavi et al. (2014a). Furthermore, the stochastic movement is slightly larger for a cell colony than for a single cell, agreeing with the results from Mousavi et al. (2014b).

The dependence of the early cell interactions and motility on the initial cell positions predicted by the model has been ascribed to the initial dominance of the mechanical signals exerted by the cells in a colony (Reinhart-King et al. 2008). However, the initial cell positions do not affect the final stage of the collective cell migration, i.e., closure of the cell-free central substrate area. This motility behaviour is qualitatively similar to experimental (Grasso et al. 2007) and numerical results on wound healing (Rey and Garcia-Aznar 2013; Vermolen et al. 2012).

Although each simulation run of the model gives a different cell position, the overall direction of the migration with random movement is the same for different simulations and consistent with *in vitro* experiments (Liang et al. 2007; Merks and Koolwijk 2009; van Oers et al. 2014; Topman et al. 2012).

It is worth noting that the predicted cell aggregation at the centre of a substrate with uniform elasticity and thickness, respectively, agrees with previous computational studies (Vermolen et al. 2015; Vermolen et al. 2012; González-Valverde and García-Aznar 2018; Chen et al. 2020) and is observed for different cell types (Palsson 2001).

### 4.3 Collective cell motility on substrates with varying elasticity and thickness

The collective cell motility on a substrate with two different elasticity regions is investigated. Whereas cells generally migrate towards stiffer substrate regions (Dokukina and Gracheva 2010; Kim et al. 2018; Mousavi et al. 2014a; Mousavi et al. 2014b), the developed model indicates that cells in a collective might not do so instantly. Instead, cells in the same substrate region initially move towards each other (at a speed depending on the substrate elasticity) to form a cluster before moving towards the substrate region with a different elasticity (Figure 6). It is argued that the mechanical signal, i.e., substrate deformation gradient, of neighbouring cells in the same substrate region is stronger than the mechanical cue from the substrate elasticity change. These results are in qualitative agreement with several studies reporting that the mechanotactic cue from the deformation of a substrate region between two neighbouring cells is stronger than the mechanical signal originating from another substrate region (Borau et al. 2011; Dudaie et al. 2015; Rey and Garcia-Aznar 2013).

The cell colonies aggregate in the substrate centre across the elasticity interface, favouring the stiff substrate region. The cell cluster originating in the soft substrate region migrates partially into the stiff substrate region. In contrast, cells in the stiff substrate region do not cross into the soft substrate region (Figure 6). The promotion of cellular interaction forces on soft substrates suggested by other studies (Guo et al. 2006; Reinhart-King et al. 2008) may explain this behaviour.

The substrate thickness variation causes a smaller difference in migration velocities of cells in the different substrate regions than the same three-fold variation in substrate elasticity. Some cells in the thin substrate region detach from the group and move to the thick substrate region (Figure 7). This is observed for cells near the thickness interface and can be ascribed to a weaker cell-cell interaction than the cue based on the thickness change, similar to reports from other studies (Tusan et al. 2018; Reinhart-King et al. 2008). Overall, cell migration is directed towards the thick substrate region. Cells migrate from the thin to the thick substrate region but not vice versa (Figure 7). This behaviour is based on a stronger mechanotactic cue from the substrate thickness change than the substrate deformation caused by neighbouring cells (Lo et al. 2000; Guo et al. 2006; Reinhart-King et al. 2008). Cell groups in the thin substrate region are attracted to the thick substrate region. In addition, the cells in a group are attached, or packed, more closely in the thick than the thin substrate region, which is observed experimentally by Tusan et al. (2018).

Collective cell migration is faster on substrates with uniform than non-uniform elasticity and thickness. The cell interactions in each of the different substrate regions increase the time required to form a final cluster in the substrate centre, i.e., decreasing the overall migration velocity.

### 4.4 Collective cell motility with cell death and division

Collective cell motility with concurrent cell death leads to a substantially longer migration period of *t* = 115 s to close a central cell-free substrate area than for the motility with cell division only and with cell death and division (*t* = 50 and 60 s, respectively) (Figure 8). The slower migration, when considering only cell death compared to the cases including cell division, is ascribed to the weaker mechanical cues. The weaker cues are associated with an 11% decrease in cell number in the case of cell death compared to increases in cell number of 60% for migration with cell division only and 47% for migration with cell division and death. However, it should be noted that the values for cell death and division probabilities in the presented simulations are exaggerated to demonstrate the effects of cell death and division.

### 4.5 Limitations

Even though the model generated results that align qualitatively with experimental data for cell migration over complex environments, the model has some limitations.

The current model considers cellular traction forces perpendicular to the substrate surface. This is inspired by the work of several groups who report that such forces do play a role in cell migration (Nerger et al. 2017; Franck et al. 2011; Maskarinec et al. 2009; Ben-David and Weihs 2021). Loof’s model based on cell forces that act normal to the substrate was used. The formalism is generic in this sense since one can also incorporate tangential forces in the same kind of formalism using a different equation for displacement in future work.

The model formalism uses a simple approach to describe cell-cell interactions mediated through the substrate. Cell-cell junctions and repulsive forces resulting from cells impinging on each other have not been considered, whereas cell deformation has been incorporated as partial overlapping. Reinhart-King et al. (2008) showed that there is a combination of repulsion and overlapping of cells. We realise that repulsion is an important process for inclusion in our future studies.

The model includes some simplifying assumptions, such as a constant cell radius, mechanical isotropy of the substrate, and the geometry of cell positions. However, these simplifications do not affect the conclusions of this work. The collective migration of cells in a multi-signalling substrate associated with different complex biological processes can be easily incorporated into the present framework. Other biological processes, such as cell differentiation (Deshpande and Spector 2017; Prokharau et al. 2014) should be considered when organ development is modelled. The nonlinearity of the problem needs an inner iteration to determine the cell positions, making an implicit method less attractive.

## Conclusion

The developed model can adequately predict collective cell motility on planar elastic substrates induced by strain energy density gradients in the substrate originating from cellular traction forces. The model considers the mechanotactic cell-cell interactions through the substrate, the substrate’s structural properties, the randomness of cell motion, and cell viability. The rate, orientation, and directionality of the cell migration depend on substrate elasticity and thickness. The model is suitable for extension to include other cell and substrates shapes and chemotactic migratory cues – increasing the potential and capabilities to replace *in vitro* and *in vivo* experiments with *in silico* simulations in, for example, wound healing, regenerative medicine, and cancer treatment.

## Statements and Declarations

### Competing Interests

Competing interests do not exist.

## Acknowledgements

The research reported in this publication is supported financially by the Organization for Women in Science for the Developing World (doctoral scholarship to RA), the European Mathematical Society (collaborative research visit award to RA), the South African Medical Research Council (grant SIR 328148 to TF), and the National Research Foundation of South Africa (grants UID92531 and CPRR14071676206 to TF). The funders had no role in study design, data collection and analysis, decision to publish, or preparation of the manuscript. Any opinion, findings, conclusions and recommendations expressed in this publication are those of the authors, and therefore the funders do not accept any liability.

## Data availability

The custom MATLAB code files of the mathematical model, supplemental videos, and data supporting this article can be accessed on the University of Cape Town’s institutional data repository (ZivaHub) under https://doi.org/10.25375/uct.17620877 as Ahmed RK, Abdalrahman T, Davies NH, Vermolen F, Franz T. Data for Mathematical Model of Mechanically Induced Collective Cell Motility on Planar Elastic Substrates, Cape Town, ZivaHub, 2022, DOI: 10.25375/uct.17620877.

## Author contribution statement

RKA contributed to Conceptualization, Data curation, Formal analysis, Funding acquisition, Investigation, Methodology, Project administration, Software, Validation, Visualization, Writing – Original Draft, and Writing - Review & Editing. TA contributed to Conceptualization, Methodology, Project administration, Supervision, and Writing - Review & Editing. NHD contributed to Conceptualization, Methodology, Supervision, and Writing - Review & Editing. FV contributed to Conceptualization, Methodology, and Writing - Review & Editing. TF contributed to Conceptualization, Funding acquisition, Methodology, Project administration, Resources, Supervision, Validation, Visualization, and Writing - Review & Editing.

## Notes

https://doi.org/10.25375/uct.17620877

### Competing Interest Statement

The authors have declared no competing interest.

